# Rare codon translation regulates growth factor-dependent cell proliferation

**DOI:** 10.1101/2025.04.17.648825

**Authors:** Yiqin Ma, Stephen M. Pronovost, Mitchell R. Lewis, Michael T. Scherzer, Jincheng Shen, Bruce A. Edgar

## Abstract

In animal cells, growth factor signaling promotes both cell cycle progression and cell growth, but the connection between these two processes is not well understood. Here, we investigated whether cell cycling and cell growth are coupled through protein translation. Using ribosome profiling and mRNA sequencing we examined changes in translational activities in human Retinal Pigment Epithelial (RPE-1) cells as they entered the cell cycle in response to serum growth factor stimulation. We found that, in addition to mRNAs encoding factors in ribosome biogenesis and translational initiation, mRNAs encoding many DNA replication factors were translationally upregulated by serum. We also noted increases of snoRNAs and tRNAs, which facilitate translation. By analyzing the distribution of 21nt mRNA ribosome footprints, produced by stalled ribosomes that lack amino-acyl-tRNAs, we found that growth factor withdrawal promoted ribosome stalling preferentially at specific rare codons. DNA replication factor genes that were translationally upregulated by serum and essential for cell cycle progression were enriched in many of these rare codons, and the cognate tRNAs that read these codons were induced by growth signaling. The serum-dependent induction of translation was more mTOR-dependent than MEK-dependent. Our results support a novel regulatory mechanism wherein growth factor signaling promotes cell proliferation by inducing tRNAs that decode rare codons, which in turn promote translational elongation of mRNAs encoding DNA replication factors to accelerate G1/S progression.

**Significance Statement:** We present a mechanistic explanation for the longstanding question of how cell cycle progression is coupled to cell growth. Our findings emphasize the role of translation machinery and reveal a “growth checkpoint” that ensures that DNA replication gene translation occurs only when growth signaling has activated the protein translation machinery. Our work complements the textbook model of the Growth Factor-Cyclin D-Rb-E2F transcriptional mechanism for cell cycle entry and is a fundamental advance for the fields of cell proliferation and growth factor signaling.

## Introduction

The proliferation of most types of cells depends on cellular growth.^1–3^ Research in many types of cells and organisms has sought to discover how cell proliferation is regulated by, coupled to, or coordinated with cellular growth, but a simple universal paradigm explaining this connection has not emerged. For animal cells, textbook explanations of growth-regulated cell proliferation focus predominantly on growth signaling via Receptor Tyrosine Kinases (RTK) and the RAS/MEK/ERK and PI3K/AKT/mTOR pathways. These signaling systems relay extra-cellular inputs in large part through cascades of protein phosphorylation and transcriptional gene induction, which culminate in the activation of G1 Cyclin/Cyclin-dependent kinase complexes (Cyclin D1/2/3-CDK4/6 and Cyclin E1/2-CDK2) and/or repression of Cyclin-dependent Kinase Inhibitors (CKIs; *e.g.* p21, p27 and p57).^1,4^ For animal cells, these models rely on the phosphorylation of retinoblastoma family proteins (pRb, p107, p130) by G1 Cyclin-CDK complexes, and/or the cell size-dependent dilution of these factors^5^, which activates E2F-type transcription factors (E2F1, 2, 3).^4,6–8^ Activating E2F activity is essential for cell cycle entry because it promotes the transcription of hundreds of cell cycle genes, including factors essential for DNA replication, mitosis, and cell cycle checkpoints.^9–11^ Although many E2F transcriptional targets are essential for cell cycle progression, the Cyclin E genes are especially important because Cyclin E-CDK2 can activate E2F by hyper-phosphorylating and inactivating Rb, leading to higher expression of Cyclin E itself and the other E2F targets. This Cyclin E-Rb-E2F positive feedback switch helps ensure that cell cycle entry is irreversible. This textbook model provides a compelling explanation of how growth signals are converted into proliferative decisions, and there is extensive support for it.^4,12–14^ However, genetic studies in model organisms indicate that this view is incomplete and perhaps overly simplistic.^1,8^ For example, mouse embryo fibroblasts lacking functional D-type or E-type Cyclin/CDK complexes can proliferate and remain growth factor dependent for cell cycle entry,^15–20^ suggesting that alternative mechanisms exist to transmit growth signals to the cell cycle machinery. Moreover, the pRb phosphorylation model doesn’t offer a specific explanation for how cell growth might be coupled to the cell cycle, apart from the fact that (as described below) growth factors also stimulate cell growth, independent of their effects on pRb, E2F and the cell cycle genes.^1^ The pRB dilution model^5^, on the other hand, does offer a mechanism for cell cycle-cell growth coupling, but still retains dependence on the activation of E2F via D– and E-Cyclin/CDK complexes.

In addition to promoting DNA replication and cell division, growth factor signaling controls growth-associated metabolism, including ribosome biogenesis and translational activity^21–26^. Growth factors also regulate metabolic processes including glucose uptake^27^, glycolysis, lactate production^28^, flux of glucose– and glutamine-derived carbons into the TCA cycle^29,30^, ribose and nucleotide biosynthesis^31,32^, mitochondrial activity^33^, and ATP production^34^. Early research showed that growth factor stimulation causes a rapid, massive increase in protein synthesis many hours before the initiation of DNA replication,^35^ and that a decrease in global translation by only 25% can delay cell cycle entry at an early stage.^36^ This indicates that the full production of certain cell cycle proteins is essential for cell division. Consistently, translational control of mRNAs encoding critical Cyclins and a CDK activator in budding yeast (Cln3) and fission yeast (Cdc13, Cdc25) can alter cell cycle rates, and cis-regulatory elements in the 5’ UTRs of these genes are responsible for this coupling of cell division to cell growth.^37,38^ Similarly, our studies in *Drosophila* found that translation of E2F1 mRNA is subject to growth signal regulation through its 5’UTR.^39,40^ Other studies in human cells revealed that mTOR signaling is essential for Cyclin D1 translation without affecting its mRNA levels.^41^ Such mechanisms could directly link growth factor signaling to core cell cycle genes via translational, as opposed to transcriptional, control. Yet although several studies suggest the involvement of translational control,^42–44^ its relative importance as a regulator of the human cell cycle is unclear, and the precise mechanisms at play remain unknown.

To address these questions, we examined growth factor-dependent translational control in non-transformed human Retinal Pigment Epithelial (RPE-1) cells using combined ribosome profiling (Ribo-Seq) and mRNA sequencing (mRNA-Seq). This method quantifies ribosome protected fragments (RPF) on mRNA and generates a genome-wide, base-pair resolution map of ribosome density for all mRNAs present in a cell.^45^ Such data is highly correlated with translational activity. We found that a large group of essential DNA replication genes are translationally upregulated by growth factor signaling. Further, detailed analyses revealed that the coding sequences of these DNA replication genes are enriched in rare codons, whose translation is selectively dependent on growth factor signaling, which induces the expression of the tRNAs that decode these rare codons.

## Results

### Identification of translationally regulated genes

To compare relative rates of transcription and translation, we conducted RNA-Seq and Ribo-Seq experiments on duplicate samples of RPE-1 cells that were first arrested in G0 by total serum withdrawal for 18 hours, and then stimulated to re-enter the cell cycle by adding 10% serum. We assayed the cells at four time points – 0, 5, 10, and 15hr after serum addition – that encompassed the cells’ transit from G0 into S phase (Figures 1A and 1B). We observed that, overall, relative fold changes of RPFs changed more dramatically than for mRNAs within each 5 hr time window, suggesting that regulation at the translational level is especially sensitive to serum growth factor signaling and probably affects the abundance of many proteins (Figures S1A-S1C). Such regulation could be important in controlling the activities of factors that mediate the quiescence-proliferation decision. Of note, temporal changes in translation were closely correlated with cell cycle status, with genes encoding factors for protein synthesis being translationally activated first (*e.g.* ribosomal protein genes, translation initiation factors), followed by DNA replication genes and then chromosome organization and mitotic cell cycle genes later on (Figures 1C-1E).

**Figure 1.**
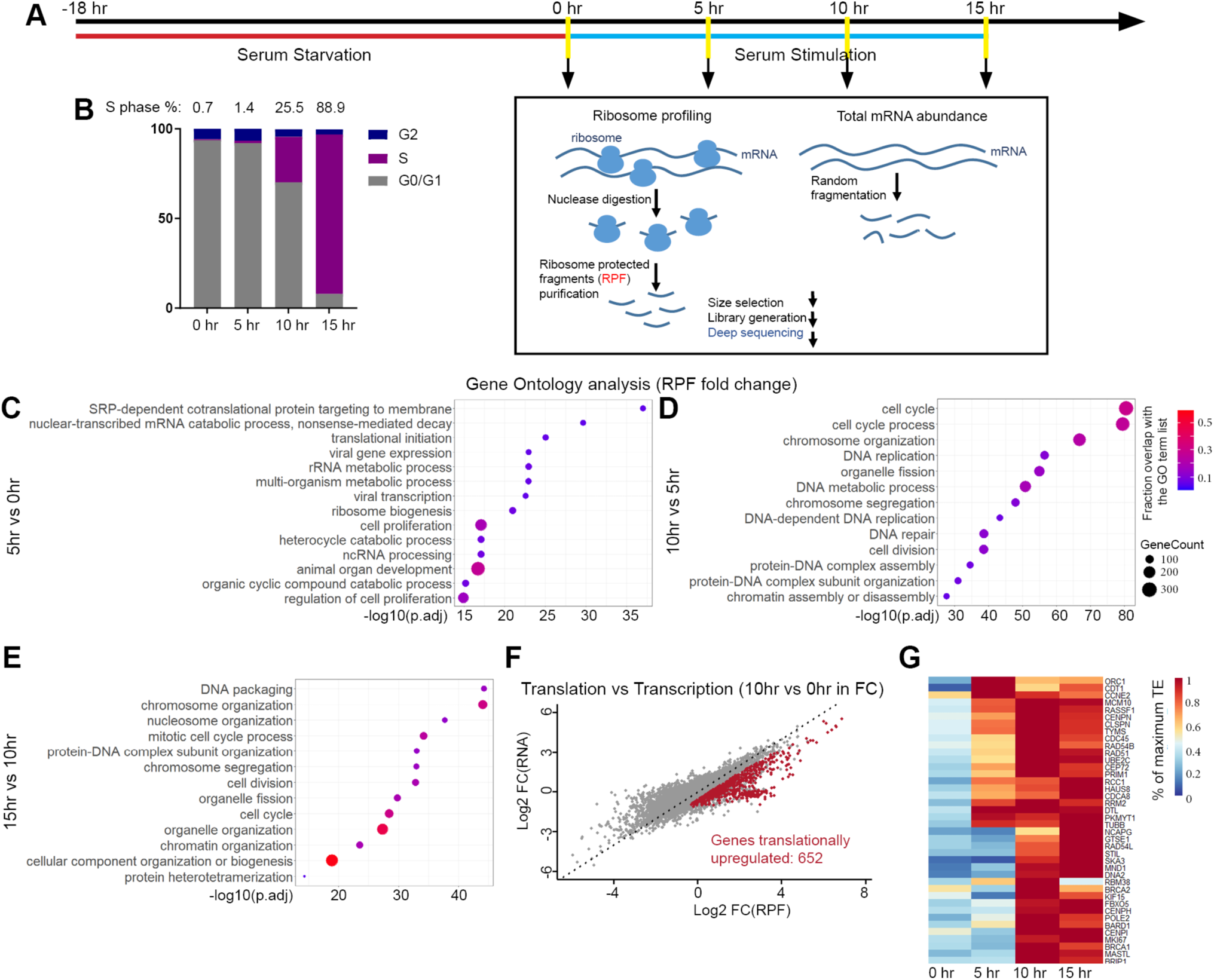
Serum-dependent translational regulation. (A) Workflow schematic of ribosome profiling and total mRNA-seq for serum-activated human RPE-1 cells. (B) Cell cycle entry kinetics of serum-stimulated RPE-1 cells, assayed by BrdU labeling and flow cytometry. (C-E) Top GO terms of genes with the highest ranked RPF fold changes between each 5 hr interval (top 1000 genes for 5hr vs 0hr and 10hr vs 5hr; all the 360 significantly changed genes for 15hr for 10hr). (F) Identification of 652 genes that are preferentially translationally upregulated, with higher RPF induction than mRNA induction, based on a 10 hr vs 0 hr comparison. (G) Heatmap of translational efficiency (TE) changes during the 15 hr serum stimulation time course, for the top 40 cell cycle genes that differed the most, based on a 10 hr vs 0 hr comparison.

Since most of the effects on cell cycle gene translation were observed within 10 hr of serum stimulation, we focused on this time-window. We identified 652 genes that were significantly upregulated for translation (*i.e.* RPF levels) relative to transcription (*i.e.* mRNA levels) by at least 1.5 fold after serum stimulation during the first 10 hr interval (Figures 1F and S1D). We observed profound enrichment in ribosomal protein genes such as RPS9, RPL15, RPS18, and RPL35, consistent with literature showing that ribosomal protein mRNA translation is growth signaling-dependent (Figures S1D and S1E).^46–48^ In addition to ribosomal protein genes, other genes involving in processes such as “cellular nitrogen compound catabolic process”, “aromatic compound catabolic process” and “DNA metabolic process” were also translationally upregulated (Figure S1D). Gene Ontology (GO) analysis of these 652 genes also revealed cell cycle genes as translationally upregulated (Figure S1D). To examine translation efficiencies, we calculated the ratios of RPF to mRNA reads that were normalized in transcripts per million kilobases (TPM). The translation efficiency of translationally upregulated cell cycle genes peaked at 5-10 hr after serum stimulation, coincident with the G1/S transition (Figures 1G, S1F and S1G). In contrast to what we had previously found in *Drosophila*,^39,40^ ribosome profiling did not show translational up-regulation of the E2F1-3 transcriptional activators (Figure S1H). We did, however, observe a large cluster of DNA replication genes including CCNE2, ORC1, CDT1, CDC6, CDC45, GINS2, POLE2, RAD51 and MCMs 2, 5, 7, and 10, that were translationally upregulated by serum stimulation (Figures 1G, S1H). These data suggest a mechanism in which DNA replication genes are regulated as a group through translation. Incidentally, many of these genes are also E2F transcriptional targets^49^, and consistent with this, our data also showed that the mRNA levels of most of these genes also rose coincident with the G1/S transition.

### Translationally regulated DNA replication genes are essential

Since not all translationally upregulated genes can be equally important for cell proliferation, we compared the 652 translationally upregulated genes with 4487 genes identified in genome-wide siRNA screens for cell cycle regulators.^50–52^ This identified 215 genes that were both translationally upregulated by serum in our analysis, and found to be required for human cell proliferation in previous studies. To identify cell cycle regulators that are rate-limiting for the quiescence to proliferation transition in RPE-1 cells, we used small interfering RNA (siRNA) suppression of these 215 genes, monitoring cell proliferation upon serum stimulation by live imaging (Figures 2A and 2B). In total, these siRNA tests identified 96 genes that are required for a normal proliferative response to serum stimulation (Figures 2C and 2D). Sixty-two of these were non-ribosomal genes (Figure 2C), with the largest group being a cluster containing 16 DNA replication and repair factors and 6 other cell cycle genes. Thus, many translationally regulated DNA replication factors are essential for normal cell cycle progression after serum stimulation.

**Figure 2.**
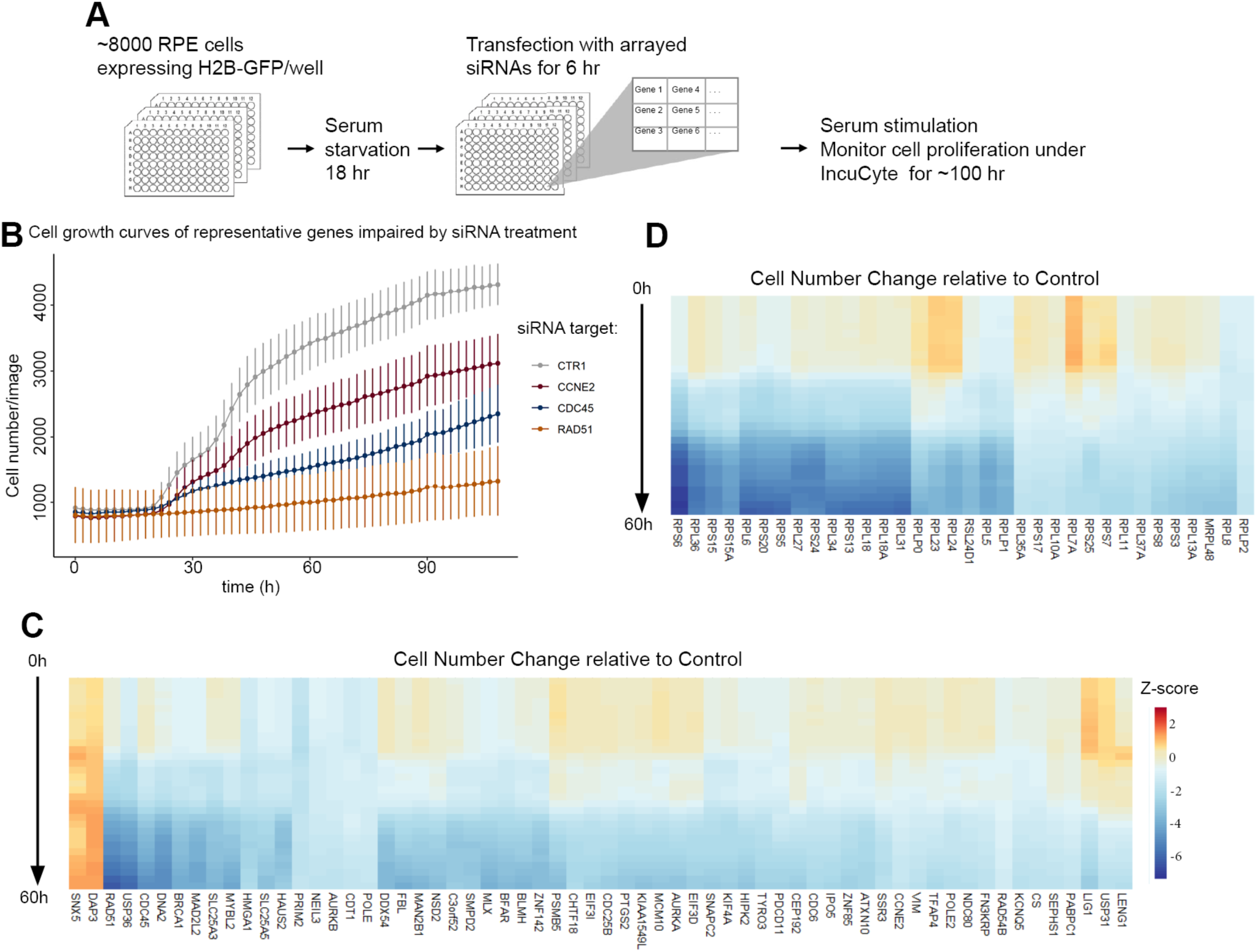
Translationally upregulated genes are required for normal cell proliferation in response to serum. (A) Schematic of siRNA screen. A total of 257 genes (gene overlap with literature was 215 plus 42 handpicked genes) were screened with 3 different siRNAs against each gene, pooled together, in 3 biological replicates. (B) Growth curves of cells during serum stimulation, treated with siRNAs against representative cell cycle genes. (C-D) Heatmap of Z-scores (number of standard deviations from control cell numbers) of non-ribosomal (C) or ribosomal (D) genes that significantly delayed cell proliferation after siRNA knockdown, during a 60h serum stimulation time course.

### Identification of Growth dependent non-coding RNAs

In addition to protein coding genes like ribosomal proteins and translation initiation factors, non-coding RNAs could also contribute to translational activity. To identify growth-dependent non-coding RNAs we carried out a microarray analysis with probes against mature transfer-RNAs (tRNAs), small nucleolar RNAs (snoRNAs), tRNA-derived small RNAs (tsRNAs) and microRNAs (miRNAs) in cells that were serum starved or stimulated for 10h (Figure 3). We detected the expression of 130 mature tRNAs, 73 of which were significantly altered by serum stimulation. The majority of these tRNAs (69 of 73) were increased after serum stimulation (Figure 3A). We also noted the upregulation of both C/D box snoRNAS and H/ACA box snoRNAs (Figure 3B), which are responsible for the 2-O-methylation and pseudouridylation of rRNAs during the rRNA maturation process.^53^ The induction of tRNAs and snoRNAs is consistent with the upregulation of translational activities by growth signaling. Our microarray also detected changes of tRNA-derived small RNAs (tsRNAs; Figure 3C-D, Figure S2). Depending on the locations derived from pre-tRNA and mature tRNA, tsRNAs can be classified into 5’Leader, tRF-1s, 5’tRFs, 3’tRFs, 5’tiRNAs, 3’tiRNAs, tRF-2s/i-tRFs. These small RNA molecules carry out diverse cellular functions such as regulating gene expression, protein translation, and cell-to-cell communication.^54,55^ Interestingly, 5’tiRNAs and 5’tRFs, but not 3’tiRNAs and 3’tRFs, have been exclusively shown to inhibit translation.^56,57^ Among the tsRNAs that were significantly downregulated by serum stimulation, we only observed 5’ tiRNAs and 5’ tRFs, and not 3’ tiRNAs or 3’ tRFs (Figure 3D, Figure S2), suggesting a possible downregulation of translation suppressors by serum. However, the exact function of the reduction in 5’tiRNAs and 5’tRFs we observed will need further characterization. The last class of non-coding RNAs we observed was miRNAs. The majority of these were significantly changed after serum stimulation were decreased (Figure 3E). GO term analysis of the genes targeted by at least 3 miRNAs that significantly decreased (top) or increased (bottom) upon serum stimulation showed primary enrichment in transcription and secondary enrichment in cell division (Figure 3F), consistent with the canonical role of miRNAs as gene expression repressors. Overall, these microarray results show that growth signaling causes widespread changes in the levels of various non-coding RNAs, many of which may stimulate translation and/or cell cycle progression.

**Figure 3.**
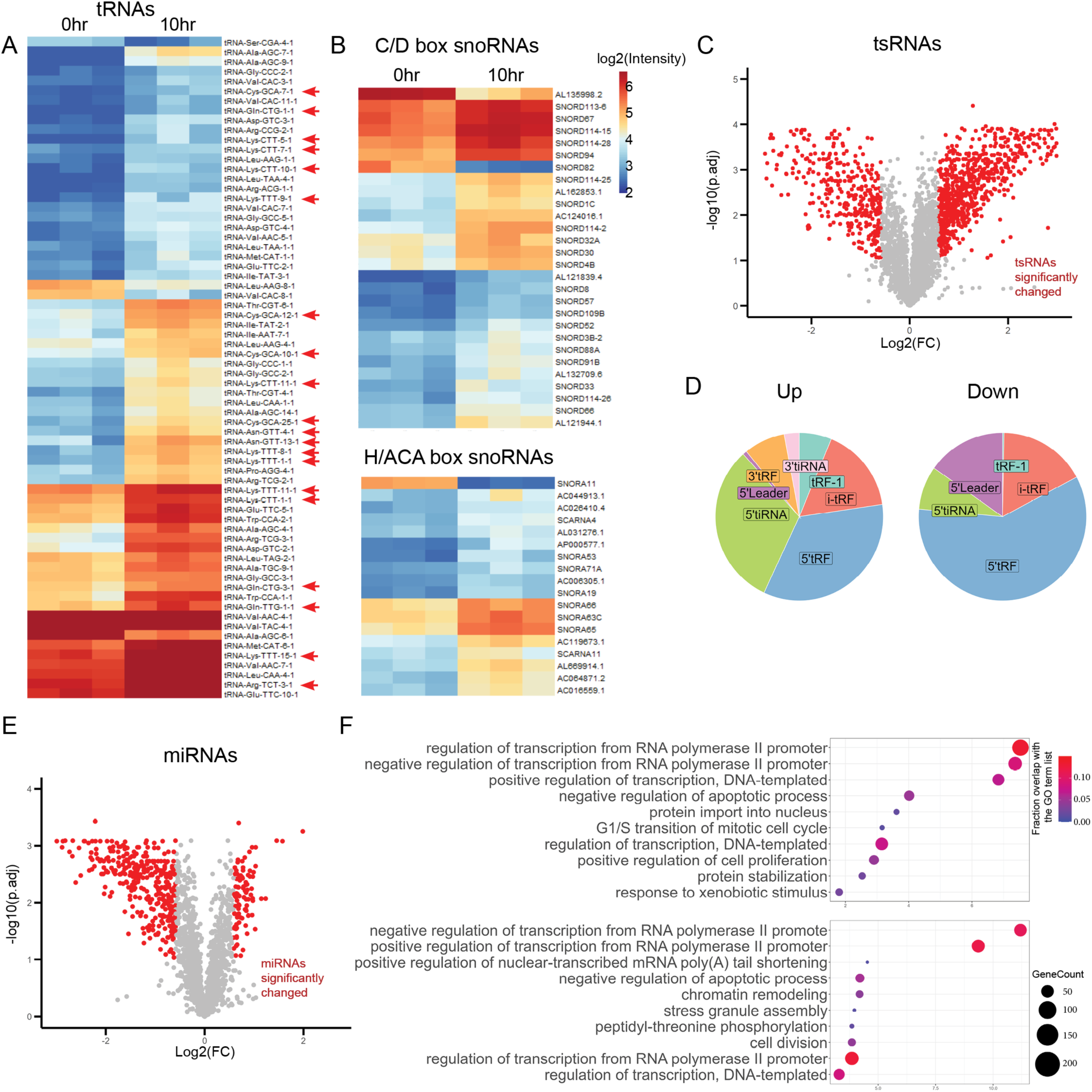
Serum-dependent regulation of non-coding RNAs. (A-B) Heatmaps of log2 normalized tRNA (A) and snoRNA (B) fold changes that are significantly regulated by serum stimulation, based on a 10 hr vs 0 hr comparison. Arrowheads in (A) indicate tRNAs corresponding to codons with high pausing indexes (Fig. 4). (C) Volcano plot of log2 fold changes vs adjusted p value for tsRNA genes. tsRNA genes with significant changes are marked in red. (D) Classification of significantly increased (left) or decreased (right) tsRNAs. (E) Volcano plot of log2 fold changes for miRNAs. miRNAs with significant changes are marked in red. (F) Top GO terms of genes that are curated in the Mirtarbase as targeted by at least 3 miRNAs that significantly decrease (top) or increase (bottom) upon serum stimulation.

### Ribosomes pause during serum starvation

Overall rates of mRNA translation are determined by the combined rates of translational initiation, elongation, and termination. Most research on translational regulatory mechanisms has focused on initiation, and data on the regulation of translation elongation is comparatively sparse. Recently, ribosomes were found to exhibit different conformations during different steps of translational elongation. Ribosomes exist in a flexible, “empty” conformation when no charged tRNA is present in the aminoacyl site (A-site). These “paused” ribosomes protect only ∼21 nucleotides (nt) of the mRNA in most nuclease-based ribosome profiling experiments^58^. In contrast, actively translating ribosomes that have a charged tRNA in the A-site span ∼28-29nt (Figure 4A).^58^ Ribosome footprints from our Ribo-Seq data indeed showed two major RPF populations of 21nt and 29nt (Figures 4A-4D, S3A and S3B). This was remarkable since our ribosome profiling protocol did not include the antibiotic, tigecycline, which stabilizes ribosomes in the 21nt conformation by blocking the initial step of codon recognition by aminoacyl-tRNAs.^58,59^ Therefore, our capture of 21 nt RFPs was sensitive even though our data probably underestimate the frequency of their occurrence. In fact, 21 nt RPF levels were >5X lower than 29 nt RPFs in all samples (Fig 4C). Both types of RPFs were concentrated in mRNA protein coding regions, and both types exhibited the same 3nt periodicity, confirming that they were true footprints of translating ribosomes on mRNAs (Figure 4D). Intriguingly, we observed a dramatic ∼4-fold decrease of 21nt RPFs relative to 29nt RPFs within 5 hrs of serum stimulation (Figure 4C). Since 21nt RPFs associate with a status of inefficient incorporation of charged tRNAs at A-sites^58^, we investigated the sequences of the corresponding A-site codons in the 21nt RPFs compared to those in 29nt RPFs. We found that the ratio of 21/29nt RPFs decreased over A-site codons coding for all amino acids after serum stimulation, albeit to varying degrees (Figure 4E, F). Overall, these results indicate that serum withdrawal causes ribosomes to stall with empty A-sites during elongation. As our tRNA expression profiling showed, this may be because charged aminoacyl-tRNAs are in short supply in serum-starved cells.

**Figure 4.**
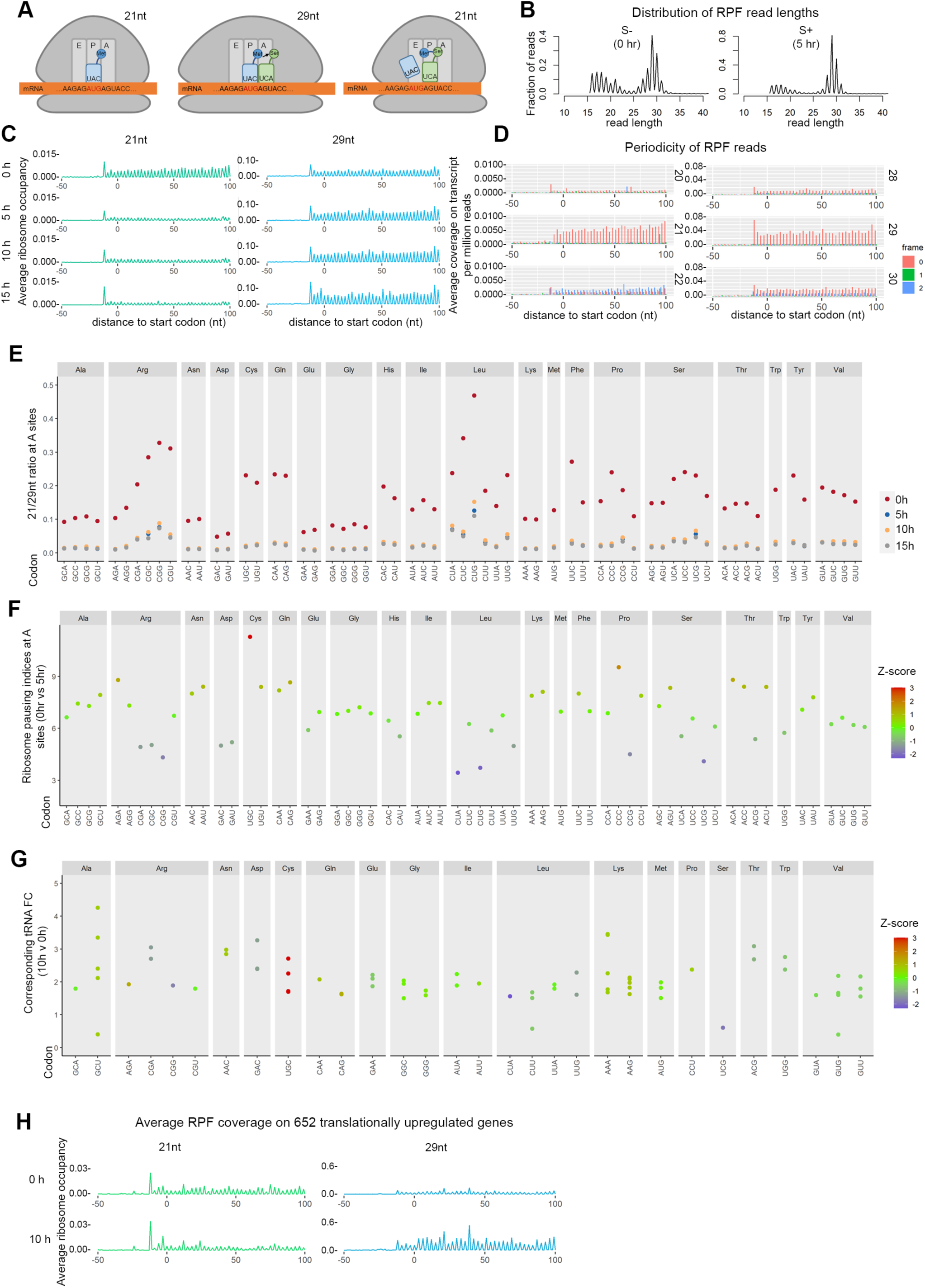
Serum stimulation reduces ribosome pausing. (A) Diagram of the translational elongation process, indicating a ribosome (grey) with A, E and P sites (light grey), an mRNA (orange), and charged tRNAs (blue, green). (B) Distribution of read lengths of ribosome protected fragments (RPFs) from serum starved cells (0 hr) and serum stimulated cells (5 hr). (C) Average ribosome occupancies on mRNA transcripts in reads per million, aligned at start codons using 21 nt RPFs (left) and 29 nt RPFs (right). (D) 3-nucleotide periodicity of different RPF read lengths aligned at start codons (average coverage on transcript per million reads) showing that 21 nt and 29 nt are *bone fide* RPFs. (E) Average ribosome pausing indices during serum starvation, at A-sites, for each codon. (F) Fold changes of significantly changed tRNAs that recognize the indicated codons, between 0 and 10hr, as assessed by microarray. Standard Z-scores for pausing indexes were presented in colors in (E-F). (G) Average ribosome occupancies of 21nt and 29nt RPFs for the 652 translationally upregulated genes, assayed at 0hr (top) and 10hr (bottom).

### tRNAs that recognize inefficiently translated codons are induced by serum growth factors

Interestingly, some codons were more enriched in 21nt RPFs than others, and so we computed a ribosome pausing index (Figure 4F) by calculating fold changes of codon enrichments in the predicted A sites in 21 vs 29nt RPFs and compared between serum starvation (0h) and stimulation (5h), for all protein coding genes. This identified several codons that had exceptionally high pausing indices at A sites, such as AGA for arginine, AAU and AAC for asparagine, UGC and UGU for cysteine, CAA and CAG for glutamine, AAG and AAA for lysine, UUC for Phenylalanine, CCC for proline, AGU for serine, and ACA, ACC, and ACU for threonine. One possible reason for unequal pausing of the A site codons is variation in tRNA abundance, so we further examined tRNA level changes from our microarray data (Figure 3A). Most tRNA genes were expressed at low or medium levels, and we observed significant serum-dependent increases of tRNAs corresponding to many of the codons with high pausing indices, including tRNA-Arg-TCT, tRNA-Asn-GTT, tRNA-Cys-GCA, tRNA-Gln-CTG, tRNA-Gln-TTG, tRNA-Lys-TTT and tRNA-Lys-CTT (Fig. 3A, 4G). The levels and translation of several mRNAs that encode tRNA synthetases also increased significantly in response to serum (*e.g.* NARS-Asn, QARS-Gln, SARS-Ser, and TARS-Thr; Figure S4A). This might increase pools of amino-acyl-tRNAs and ameliorate 21-mer RPF pausing. Likewise, we observed strong increases in mRNAs and RPFs that mapped to tRNA modification enzymes,^60^ suggesting a third potential mechanism for differential ribosome pausing during serum starvation (Figure S4B). Based on the likelihood that 21nt and 29nt RPFs indicate paused versus elongating ribosomes, we then examined the distribution of 21 vs. 29nt RPFs in our set of 652 serum-dependent translationally upregulated genes. On average we found strong increases in 29nt RPFs and moderate decreases in 21nt RPFs spanning the coding regions of these 652 genes after 10 hr serum stimulation (Figures 4F and S3C). This correlation further supports our proposal that the presence of 29nt RPFs indicates a higher efficiency of translation.

### DNA replication genes are enriched in rare codons

A previous study reported that functionally related genes share similar codon usages, with cell cycle genes and cell adhesion and patterning genes exhibiting distinct profiles.^61^ To investigate whether certain codon preferences are correlated with higher translation rates after serum stimulation, we examined the codon usage among different gene groups (Figures 5A, S5A). Interestingly, the set of translationally upregulated DNA replication genes that we had defined has a codon usage profile distinct from that of our entire set of 652 translationally controlled genes, or from the ribosomal genes that are also serum-responsive for translation. This profile shows over-representation of codons that are rarely used in protein coding genes at large, including codons with high pausing indexes such as AAU for asparagine, UGU for cysteine, AGU for serine, and ACA for threonine, *etc.* (Figures 5A and S5A). This pattern is consistent with a report showing that mRNAs which oscillate during the cell cycle are enriched in rare codons and undergo a translation boost upon serum stimulation.^62^ The levels of tRNAs for rare codons are typically low,^63^ and therefore ribosome pausing at these rare codons may become rate limiting for translation elongation, constituting an unusual post-transcriptional mechanism for controlling gene expression that could be serum-responsive.

**Figure 5.**
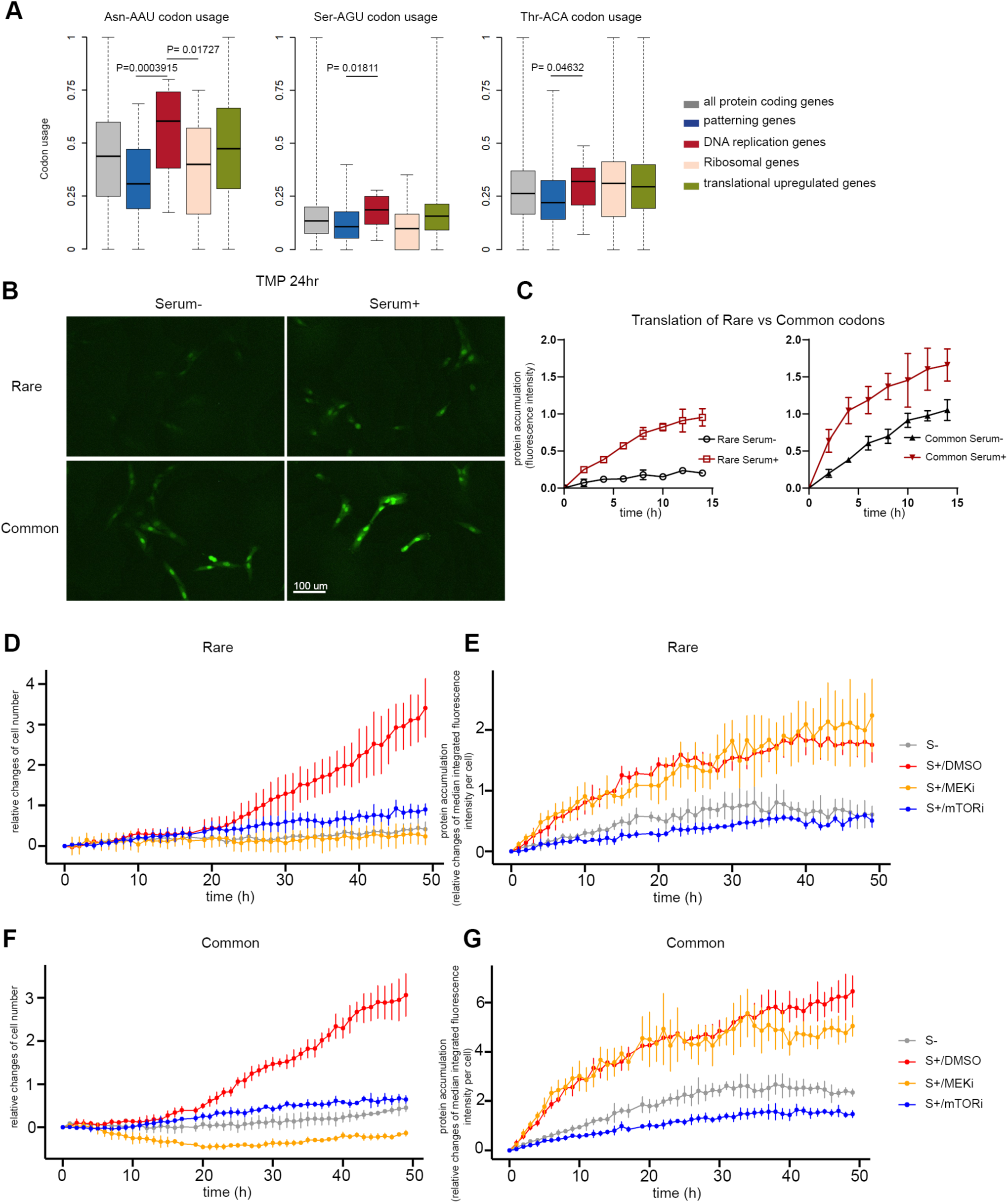
Rare codon usage safeguards translation of DNA replication genes. (A) Usage of the example rare codons Asn-AAU, Ser-AGU and Thr-ACA in protein coding genes, patterning genes, DNA replication genes from our siRNA screen, ribosomal genes, and the 652 translationally upregulated genes. P-values were determined by Wilcox tests. (B) Fluorescent images of Rare and Common reporters under serum starvation or stimulation for 24h. (C) Relative increase of fluorescence intensity for Rare codon and Common codon reporters under serum starvation or serum stimulation. Data are the mean (±1 SD). P-value=0.0006 (for slopes of Rare S+ vs Rare S-). P-value=0.0544 (for slopes of Common S+ vs Common S-), two-tailed t tests. (D-G) Relative changes of cell number (D, F) or median integrated GFP fluorescence intensity per cell (E, G) for the Rare codon and Common codon reporters, under serum starvation (S-) or serum stimulation (S+), and treated with inhibitors against MEK (MEKi), mTOR (mTORi), or control (DMSO, solvent). Data are the mean (±1 SD).

### Rare codon translation is especially serum-dependent

To further determine the impact of codon usage on translation, we constructed codon reporter genes in which the coding region of an sfGFP-ecDHFR fusion protein uses codons that are relatively rare and preferentially present in DNA replication genes, or, alternately, the codons that are most common in all genes, without changing the amino acid sequence (Figure S6A). ecDHFR is a controllable degron, such that addition of its ligand trimethoprim (TMP) stabilizes the otherwise unstable sfGFP-ecDFHR fusion protein. When this reporter is used, the ensuing accumulation of GFP fluorescence after TMP addition directly mirrors the translation rate of the test mRNA.^64^ We found that the translation efficiency of the reporter enriched in common codons was high even without serum stimulation, and that it increased <2-fold upon serum stimulation. In contrast, the reporter enriched in rarer, DNA replication gene-associated codons was nearly silent in the absence of serum and responded strongly to serum, increasing its rate of translation nearly 5-fold (Figures 5B,C, S6B). This result suggests that changes in codon usage can function as a highly sensitive growth-signaling detector.

### mTOR, but not MEK, is required for serum-dependent translational control on codon usage

Serum stimulation activates multiple growth signaling pathways, among which RAS/MEK/ERK and PI3K/AKT/mTOR signaling are the major pathways known to directly regulate tRNA biogenesis and protein translation.^65–69^ To test their importance in regulating DNA replication gene translation, we treated cells with either a MEK inhibitor (Trametinib) or an mTOR inhibitor (GDC-0349) during serum stimulation and then assayed translation using our reporters. We found that only the mTOR inhibitor was able to suppress the translation of the sfGFP-ecDFHR translation reporters (Figure 5E, 5G), despite that both inhibitors strongly arrested cell proliferation (Figure 5D, 5F). Notably, mTOR inhibition suppressed translation from both the rare-codon and common-codon reporters approximately equally, indicating that mTOR does not selectively affect translation in a codon-dependent manner.

## Discussion

The dependence of cell division on cell growth was first described more than a century ago.^70^ To explain this dependence, existing paradigms of growth-dependent cell proliferation have emphasized the growth factor-dependent transcriptional activation of cell cycle genes.^1–4^ In this study we describe a different mechanism in which translational activity regulates the cell cycle. Because cell growth also requires the mobilization of translation^35,36^ this mechanism can couple S-phase entry to a cell’s growth status quite directly, and (in principle) independently of transcriptional control mechanisms.

We found that a large set of DNA replication genes are translationally upregulated by serum-dependent signaling, and that these genes are enriched in a special set of codons that are under-represented in other classes of genes. In reporter gene tests, these codons throttled translation nearly completely in the absence of serum but allowed rapid translation after serum stimulation. In contrast, a matched reporter gene with common codon usage was efficiently translated even during serum starvation. These disparate effects suggest the existence of a translationally mediated “growth checkpoint” that preferentially operates on genes enriched in specific rare codons, *e.g.* DNA replication genes. We propose that this checkpoint surveils growth factor signaling activity and can act as a safeguard to suppress the expression of DNA replication proteins, and S-phase entry, if cells activate transcription of the DNA replication genes in non-permissive conditions. This checkpoint might be engaged, for instance, in cancer cells that have Rb mutations and high E2F activity but grow in a tumor microenvironment environment that is poor in nutrients or growth factors. The enrichment of rare codons in DNA replication genes may also provide a convenient mechanism for rapidly expressing DNA replication factors *en masse* in response to growth signaling, and prior to their full transcriptional activation in response to rising E2F activity. This could be necessary for a coordinated and timely entry into S phase. Our results are partially in agreement with a report on murine cells^62^ which found that dynamic mRNAs whose expression increases during each cell cycle are enriched in rare codons. That study also presented a rare codon translational reporter gene that exhibited a stronger translational boost in high serum than a matched common codon reporter. Though some details diverge, both that study^62^ and ours indicate conserved translational regulation via rare codons.

tRNA availability is crucial for efficient translation. Our data are consistent with a mechanism in which growth signaling promotes translational activities in part by upregulating the expression of a select set of tRNAs. Our results showing tRNA inductions in response to serum (Fig. 3) are consistent with other reports showing that tRNA expression profiles are subject to extensive regulation based on proliferative or metastatic status.^61,71–75^ When the availability of tRNAs is low, ribosomes tend to pause on mRNAs with empty A sites, which manifest as 21nt ribosome protected fragments in Ribo-Seq data. This pausing effect was especially obvious for the rare codons we identified. However, when growth signaling is activated, tRNAs, especially those that recognize rare codons, are induced. The induction of tRNAs quickly releases ribosomes from pausing and boosts translation efficiency. The serum-dependent induction other translation promoting factors, such as tRNA sythetases (Fig. S4A), snoRNAs (Fig. 3B), tsRNAs (Fig. 3C), and the repression of translation-supressing miRNAs (Fig. 3E), may also contribute to the translational upregulation of DNA replication genes. Our results further indicated that translational control is highly dependent on the mTOR signaling pathway, and less sensitive to RAS/MEK/ERK signaling (Fig. 5). However, the mechanism that makes rare codon translation selectively serum-dependent is likely to be both MEK– and mTOR-independent, raising a puzzle for future investigations.

In summary, our study reveals a sensitive, somewhat novel mechanism for growth-signaling dependent cell cycle control (Fig. 6). In this mechanism, serum growth factors induce tRNAs and other factors that jump-start the translational elongation of mRNAs, especially those enriched in rare codons that are common in DNA replication genes. Efficient translation in the presence of serum then ensures the timely accumulation of DNA replication proteins and S phase entry. In the absence of serum or other factors necessary for cell growth (*e.g.* amino acids) this mechanism may constitute a “growth checkpoint” that delays S-phase onset, thereby coordinating cell cycle progression with protein synthesis-dependent cell growth. In principle, codon usage, tRNA gene expression, and the expression of tRNA synthetases and modification enzymes offer expansive domains that may have been tuned during evolution to alter gene expression programs. We suggest that a thorough understanding of translational elongation and tRNA metabolism will be required for a comprehensive understanding of not only cell cycle control, but other important biological processes.

**Figure 6.**
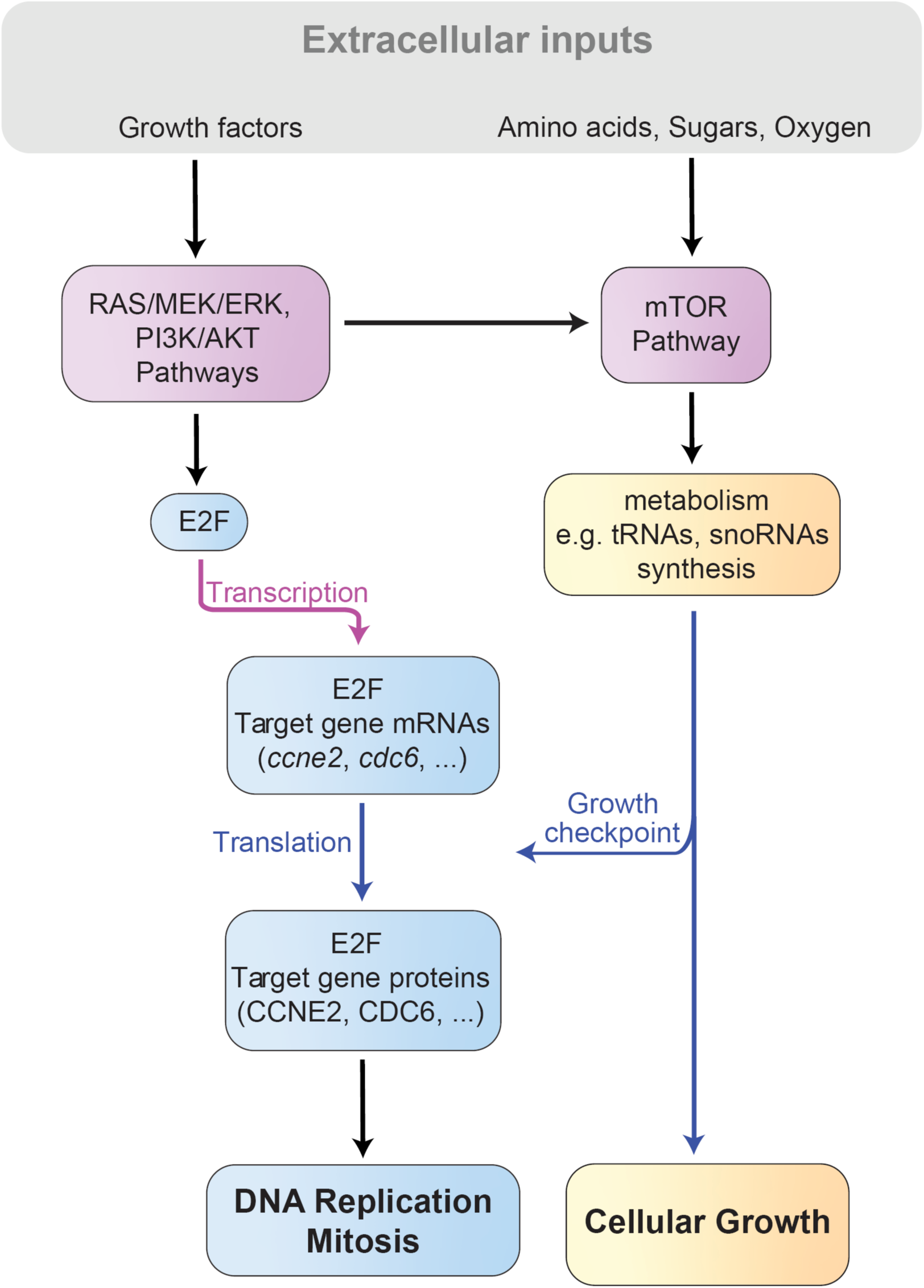
Model diagraming how growth factor signaling regulates the cell cycle by regulating the translation of cell cycle gene mRNAs.

## Materials and Methods

### Materials

Human RPE-1 cells were obtained from ATCC and cultured in Complete Medium (DMEM/F12 with 10% FBS, L-Glutamine and Penicillin-Streptomycin (all Gibco) at 37 °C and 5% CO_2_. For serum starvation experiments, 4×10^5^ cells per 10cm plate were plated and grown for 24 hours in Complete Medium prior to starvation. Cultures were washed with 1xPBS, and then incubated for 18-24hr in Starvation Medium (DMEM/F12 with L-Glutamine and Pen-Strep). To initiate serum stimulation, Starvation Medium was replaced by fresh Complete Medium.

For serum stimulation assays with inhibitors, DMSO, Trametinib (MEK inhibitor, working concentration 100 nM), GDC-0941 (PI3K inhibitor, working concentration 5 µM) or GDC-0349 (mTOR inhibitor, working concentration 1 µM) were added to the medium 1hr before PBS wash and serum stimulation. Then cells were incubated with Complete Medium mixed with the corresponding inhibitors or equivalent amount of DMSO (the solvent for inhibitors).

293FT cells were obtained from the Mary Beckerle Lab (Huntsman Cancer Institute, University of Utah, Salt Lake City, USA) and were cultured in DMEM, supplemented with 10% FBS, 1× L-Glutamine, 1x Sodium Pyruvate and 1× MEM NEAA (all Gibco) at 37 °C and 5% CO2.

## Methods

### Ribosome profiling and total RNA-seq sample collection

Cell cultures were washed with 1xPBS prior to flash-freezing in ethanol/dry ice and storage at – 80C°. Ribosome profiling samples were quickly thawed in parallel, and immediately lysed for 10 minutes on ice in 400 μl/plate of cold Lysis Buffer (10mM Tris –HCl pH 7.5, 300mM KCl, 10mM MgCl2, 1% Triton X-100, 1mM DTT, 4U DNase, 0.2mg/ml cyclohexamide). Cell lysates were collected by scraping, then passaged 10x through a 26G° needle prior to centrifugation for 10 min at 17,000x g, 4°C. RNase1 (750U/sample, Thermo Fisher) was added and incubated for 45 minutes at RT before RNase digestion was stopped by the addition of 200U/sample SUPERaseIn (Invitrogen). Samples were vortexed gently to mix before centrifuging 12,000x g for 10 min at 4°C. Supernatants were split into 220 µL aliquots and underlaid in 11×34 mm polycarbonate tubes (Beckman) with 660 µl of Sucrose Buffer (10mM Tris –HCl pH 7.5, 300mM KCl, 10mM MgCl2, 50% Sucrose, 13.2 U SUPERaseIn, 3.3 mM MgCl2, 0.5 mM DTT, 0.2mg/ml Cyclohexamide) before centrifuging at 200,00x g at 4°C for 3.5 hours in a TL-100 rotor/ ultracentrifuge (Beckman). Supernatants were removed and ribosomal pellets from each sample were resuspended in a combined volume of 700 µL Qiazol (Qiagen) for RNA isolation with miRNeasy Mini kits (Qiagen).

For total RNA-seq sample collection, parallel cultures were thawed and lysed in 700 µl/sample Qiazol (Qiagen). Lysates were collected by scraping and total RNA was purified via miRNeasy Mini kit (Qiagen). Both Ribo-seq and RNA-seq samples were carried out with 4 biological replicates.

### Library construction

rRNA was depleted using Ribo-Zero Gold rRNA Removal Kit (Illumina, discontinued). Ribo-Zero treated total RNA and RPFs were purified using RNA Clean and Concentrator-5 kit (Zymo). Before library construction, total RNA samples were heat fragmented. Briefly, 7 µl of each Ribo-Zero treated total RNA sample was applied with 3 µl of 10x T4 Polynucleotide Kinase Buffer (700 mM Tris-HCl pH 7.6, 100 mM MgCl2, 5 mM DTT) and mixed on a Eppendorf Thermomixer at 2000 rpm for 30 sec before transferring to a thermal cycler with a heated lid set at 100°C and heated at 94 °C for 8 min, followed by ice bath for 1 min. Both RPFs and fragmented RNAs were then treated by T4 Polynucleotide Kinase in T4 Polynucleotide Kinase Buffer supplemented with 1.2 mM ATP, 24 U T4 Polynucleotide Kinase and 40 U RNaseOUT (ThermoFisher Scientific) per 50 µL reaction at 37 °C for 60 min before purification using RNA Clean and Concentrator-5 kit (Zymo). Libraries were constructed using Qiagen QIAseq miRNA library kit by Dr. Brian Dalley, High-Throughput Genomics Shared Resource, Huntsman Cancer Institute, University of Utah.

### Sequencing

Libraries were sequenced on an Illumina HiSeq 2500 instrument with 50 cycle single-read.

### Data analysis

Reference genomes and annotations were obtained from Ensembl release 90 (Homo_sapiens.GRCh38). Adapter sequences were removed and UMI tags were moved into the FASTQ header by a customized PerL script from Dr. Chris Stubben and Dr. Timothy Parnell, Bioinformatic Analysis Shared Resource, Huntsman Cancer Institute, University of Utah. Both ribosome profiling and RNA-seq reads were aligned to the reference genome using STAR^76^ version 2.5.4a with the following parameters: –-outFilterMismatchNoverLmax 0.05 –-outFilterMatchNmin 16 –-outFilterScoreMinOverLread 0 –-outFilterMatchNminOverLread 0 –-outFilterMultimapNmax 50 –-winAnchorMultimapNmax 200 –-seedSearchStartLmax 13 –-alignIntronMax 1 –-outWigType bedGraph –-outWigStrand Unstranded. Duplicates with the same UMI tag and position were marked with a customized PerL script. In order to examine the truthful translational activities without being confounded by translation initiation and termination, we only focused on the core CDS region from the 15th codon through the 6th to the last codon for each transcript. Altered Gene transfer format (GTF) files with a focus on the core CDS regions were generated in R version 3.5.1 using customized script. Uniquely mapped, non-duplicated reads were counted using featureCounts version 1.5.1 with parameters –T 8 –s 1 –-largestOverlap –-ignoreDup –t ‘CDS’.

Differential expression analysis was carried out in R using DESeq2^77^ version 1.20.0. Significance of RNA and RPF expression changes were determined by at least 2 fold change and padj < 0.05. Only reads that uniquely mapped to a single genomic region were used. Translationally regulated genes were determined by log2FC difference between RPF and RNA larger than 0.58 and padj < 0.1 from xtail package^78^ version 1.1.5. Metagene and codon analysis were carried out with customized Rscripts. GO term analysis was carried out with DAVID^79^ and terms were summarized using Revigo.^80^ Heatmaps were generated using R package pheatmap version 1.0.10. Scatterplots, dot plots and box plots were generated using ggplot2 version 3.3.6.

### Pausing Index calculations

Per codon Ribosome Pausing Indexes were calculated as dividing the ratio of 21/29 nt at 0hr by the ratio of 21/29nt at 5hr=Ratio_0hr_ ^21/29 nt^/ Ratio_5hr_ ^21/29 nt^. To avoid translation initiation and termination as possible confounders, we used only core CDS regions, starting from the 15th codon and running through the 6th to the last codon for each transcript for all protein coding genes. We consistently observed a 12 nt P-site offset (the distance from the 5’ end of the read to the ribosomal P-site) for both 21 nt and 29 nt RPFs (Figure 4C). Therefore, we determined A-site offsets and E-site offset as 15 nt and 9nt, respectively.

### siRNA screen

siRNA library against selected 257 translationally upregulated genes was purchased from Bioneer in the format of 3 siRNAs pooled (0.1 nmole each) for each gene in 96 well plates. Each plate also contained siRNAs against PLK1 and CCNE2 as positive controls, and luciferase GL2 and a negative siRNA from Bioneer as negative controls. Utilizing a Tecan EVO100/MCA96 Liquid Handler in sterile bio-hoods from the Drug Discovery core of University of Utah, 8000 RPE-1 cells expressing H2B-GFP were placed into each well of 96 well plate (Corning) in 100 µL complete medium. After settling for 24h, cells were washed with 1xPBS and then incubated in 100 µL DMEM/F12. After 18h starvation, pre-mixed siRNA transfection complexes containing 10 µL Opti-MEM (Gibco), 0.44 µL of 2 µM siRNAs, 0.3 µL TransIT-X2 (Mirus Bio) were added to corresponding wells. After 6h incubation (a total of 24h serum starvation), cells were released from serum withdrawal by exchanging medium to fresh 100 µL complete medium. Then plates were transferred into incuCyte (Sartorius) to monitor the real-time cell growth at every 2 hr for 120 hrs in total. 3 biological replicate plates were treated and monitored in parallel. Cell numbers were determined and counted based on H2B-GFP signals by the analysis tool of incuCyte. Significance of growth curve changes compared to control was determined by simulations using statmod package in R.

### Microarray of non-coding RNAs

3 replicates of 0h and 10h total RNA-seq samples were applied to Arraystar small RNA microarray. Specifically, the RNA samples were quantified by NanoDrop ND-1000 spectrophotometer and RNA integrity examined by Bioanalyzer 2100 or gel electrophoresis. For each sample, 100 ng total RNA was firstly dephosphorylated to form the 3-OH end. The 3-OH ended RNA was then denatured by DMSO and enzymatically labeled with Cy3. The labeled RNA was hybridized onto Arraystar Human small RNA Microarray (8×15K, Arraystar) and the array was scanned by an Agilent Scanner G2505C. Agilent Feature Extraction software (version 11.0.1.1) was used to analyze acquired array images. Quantile normalization and subsequent data processing were performed using GeneSpring GX v12.1 software package (Agilent Technologies). After normalization, the probe signals having Present (P) or Marginal (M) QC flags in at least 3 out of 6 samples were retained. Multiple probes from the same small RNA (miRNA/TsRNA (tRF&tiRNA)/pre-miRNA/tRNA/snoRNA) were combined into one RNA level. Differentially expressed small RNAs between two comparison groups were identified by fold change (FC >1.5) and statistical significance (p-value <0.05) thresholds.

### Translation Codon reporters

The CDS region from the 15th codon till last 6th codon of NLS-sfGFP-NLS-ecDHFR fusion protein was altered as in Figure S6A. DNA fragments were commercially synthesized from Geneuniversal and cloned into the pHIV-Zsgreen vector using XbaI and ClaI restriction enzyme sites. Vectors were sequenced for validation by the DNA Sequencing Core, University of Utah. Lentiviruses were generated by transfecting 80-90% confluent 293FT cells in 6-cm plates with 500 µL Opti-MEM (Gibco), 3.75 µg pHIV-Zsgreen plasmid, 2.82 µg psPAX2 plasmid, 0.93 µg pMD2.G plasmid, 15 µL TransIT-X2 (Mirus Bio) for 24 hr before changing medium. 36-48 hr later, medium containing virus was collected and filtered using cellulose acetate 0.45 µm sterile filter (Corning), and then applied at 1:1 ratio to each well of 6-well plate containing 0.4 x 10^5^ RPE-1 cells in complete medium supplemented with 8 µg Polybrene (Millipore). 24hr after virus incubation, medium was replaced with 2ml fresh complete medium. RPE-1 cells successfully expressing high copies of codon reporters were sorted by Aria 5 Laser Cell Sorter (BD) in the Flow Cytometry Core, University of Utah. 800 cells expressing corresponding codon reporters were placed into 96 well plate for settlement. Then cells were serum starved for 24h before they were reintroduced to complete media supplemented with serum and containing DMSO (solvent) or 10 µM TMP (Cayman Chemical). GFP fluorescence was monitored at every 2 hr in an IncuCyte or LiveCyte (Phasefocus) instrument.

### Generating stable RPE-1 stable cell lines that expressed H2B-GFP

Lentiviruses bearing H2B-GFP were generated by transfecting 80-90% confluent 293FT cells in 6-cm plate with 500 µL Opti-MEM (Gibco), 3.75 µg PGK-H2BeGFP plasmid, 2.82 µg psPAX2 plasmid, 0.93 µg pMD2.G plasmid, 15 µL TransIT-X2 (Mirus Bio) for 24 hr before changing medium. 36-48 hr later, medium containing virus was collected and filtered using cellulose acetate 0.45 µm sterile filter (Corning), then was applied at 1:1 ratio to each well of 6-well plate containing 0.4 x 10^5^ RPE-1 cells in complete medium with 8 µg Polybrene (Millipore). 24hr after virus incubation, medium was replaced with 2ml fresh complete medium. RPE-1 cells successfully expressing high copies of H2B-GFP were sorted by Aria 5 Laser Cell Sorter (BD) in the Flow Cytometry Core, University of Utah.

### Quantification of codon reporter’s genomic insertions and mRNA expression

Genomic DNA of RPE-1 cells expressing rare or common codon reporters was isolated using QIAamp DNA mini kit (Qiagen).

Total mRNA from serum starved or serum stimulated cells were isolated using RNAqueous-Micro Kit (Invitrogen). First strand cDNA was synthesized by ProtoScript II Reverse Transcriptase (NEB).

Rare codon genomic DNA or cDNA was detected by primers: Fwd-GGAATCAATCGGTCGTCCTT, Rev-TGGCTTCATCCACACTCTTC. Common codon genomic DNA or cDNA was detected by primers: Fwd-CACCCAAAGCGTTCTGTCTA, Rev-TTTGTACAGCTCGTCCATACC. CCNE2 3UTR served as genomic DNA reference control and was detected by primers: Fwd-GAAGCCAACCACAGTCTATACC, Rev-GGTAGCACTTATCCAGTCCAAA. mRNA quantification for CCND3 was detected by primers: Fwd-TGGCTTACTGGATGCTGGAGGTAT, Rev-ACAGGTAGCGATCCAGGTAGTTCA; Cdt1 was detected by Fwd-GAAGCTGGTGGAGCATGTCA, Rev-TGTCGGGTACTTCATCCACG; CCNE2 was detected by Fwd-CGGCCTATATATTGGGTTGGCG, Rev-ACGGCTACTTCGTCTTGACATT. Housekeeping gene RPS20 cDNA served as internal control and was detected by primers: Fwd-GCGACTCATTGACTTGCACA, Rev-TCAAAGTGTACTGCTGGCCC.

q-PCR reactions were carried out using iTaq Universal SYBR Green Supermix (Biorad) in a CFX384 Touch Real-Time PCR Detection System (Biorad).

### BrdU incorporation and FACS

RPE-1 cells were incubated with 10 µM BrdU in 10 cm plates for 30 min before PBS wash and dissociation by 0.05% Trypsin (ThermoFisher). Trypsinization was terminated by adding complete medium to the plates and cells were precipitated by centrifuge at 1500 rpm, 5 min. Cell pellets were thoroughly washed in PBS and fixed in cold 70% EtOH. Cells were then acidified and permeabilized by 2N HCl, 0.5% Triton-X100 for 30 minutes at RT, followed by neutralization with 0.1M Na Borate pH 8.5. After PBS wash, cells were subjected to immunostaining with primary antibody mouse anti-BRDU (BD Clone B44) at 4C overnight and secondary antibody goat anti-mouse Ig Alexa 488 (Invitrogen, A-11001) for 30 min at room temperature in PBS containing 1% BSA, 0.5% Triton. Finally, DNA was stained with 5 µg/µL Propidium Iodide before the cells were examined using a Cytoflex LX.

## Data Availability

All RNA sequence data used in this study, including RNA-Seq and Ribo-Seq data are available in the GEO public database under the accession number GSE227197^81^. All other raw data that are not included in this paper’s supporting information files are available upon request from the authors. Custom RScripts used in this analysis are available upon request from the authors. All other code is publicly available.

## Acknowledgments

We thank Michael Howard for ribosome profiling protocols, Hanlin Zeng, Chris Jensen, Mona Foth and Martin McMahon for reagents, Ozlen Balcioglu and Robert Judson-Torres for IncuCyte and LiveCyte imaging assistance, and the Huntsman Cancer Institute High Throughput Genomics Core Facility (Brian Dalley) and Bioinformatic Analysis Shared Resource for sequencing and data analysis assistance. We also thank the University of Utah Health Sciences Center DNA/Peptide Synthesis Core Facility, DNA Sequencing Core Facility, Drug Discovery Core Facility and Flow Cytometry Core Facility for their services. This work was supported by NIH grants GM126033 and GM140900 to BAE, P30CA042014 to the Huntsman Cancer Institute, and S10 OD021644 to the Center for High Performance Computing (CHPC) at University of Utah.

## Author Contributions

Conceptualization: YM, SMP, BAE

Methodology: YM, SMP, MRL, BAE

Software: YM, MRL, JS

Validation: YM

Formal analysis: YM, MRL, JS

Investigation: YM, SMP, MRL, MTS, EW

Resources: BAE

Data Curation: YM

Visualization: YM

Funding acquisition: BAE

Project Administration: BAE

Supervision: BAE

Writing – original draft: YM, BAE

Writing – review & editing: YM, SMP, MRL, BAE

## Classification

Major classification: Biological Sciences; Minor Classification: Cell Biology

## Figures and Tables

**Figure S1.**
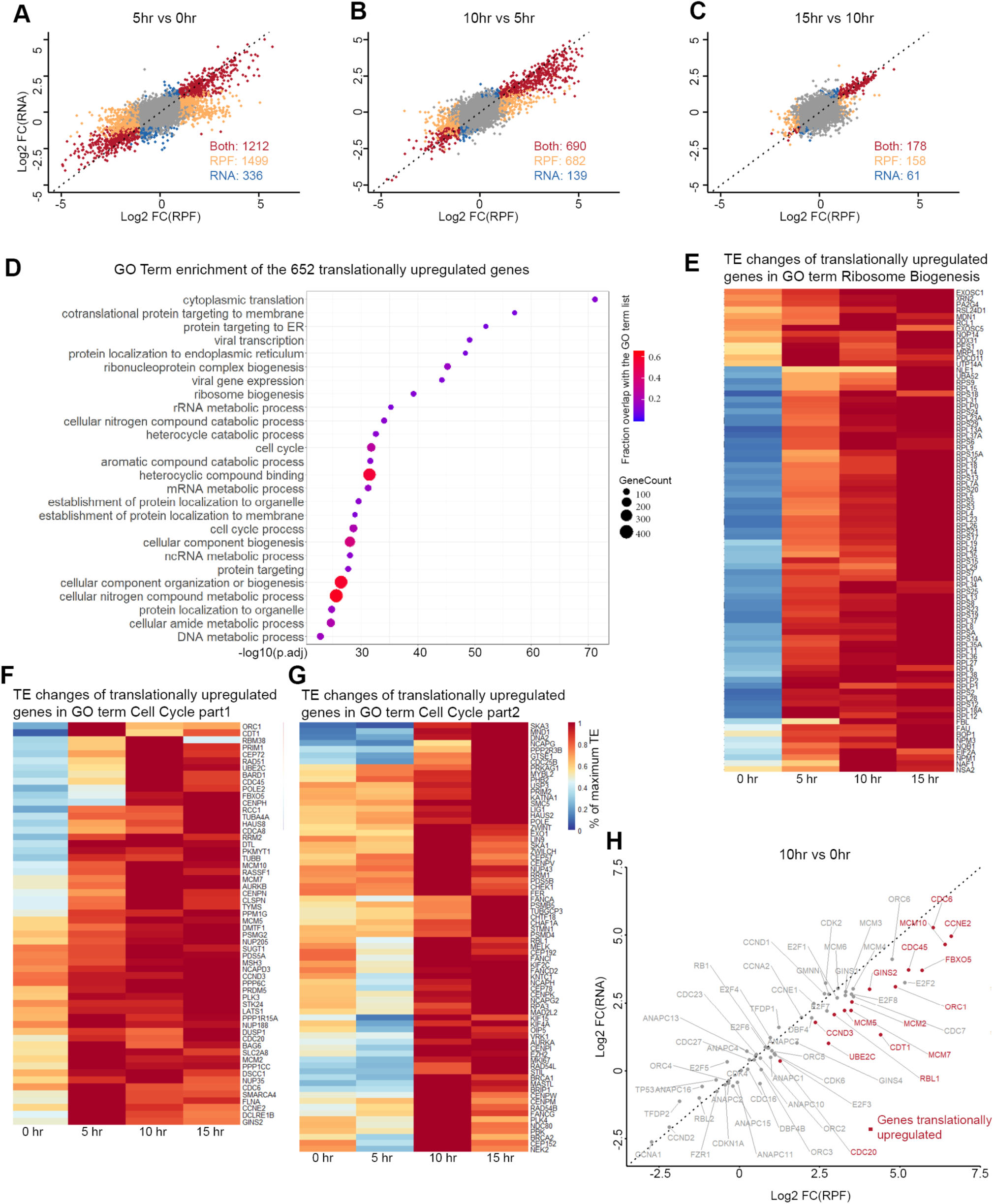
Ribosomal protein and cell cycle genes are translationally upregulated by serum growth factors. (A-C) Log2 fold changes of mRNA vs ribosome protected fragment (RPF) abundance between each 5 hr interval (5hr vs 0hr, 10hr vs 5hr, 15hr vs 10hr) after serum stimulation. (D) Top GO terms of the 652 genes that were differentially upregulated in translation between 10 hr and 0 hr. (E-G) Heatmaps of translation efficiencies (TE) during the 15 hr serum stimulation time course for ribosomal genes (E) and cell cycle genes (F-G). (H) RNA vs RPF log2 fold changes of selected key cell cycle regulators between 10hr and 0 hr.

**Figure S2.**
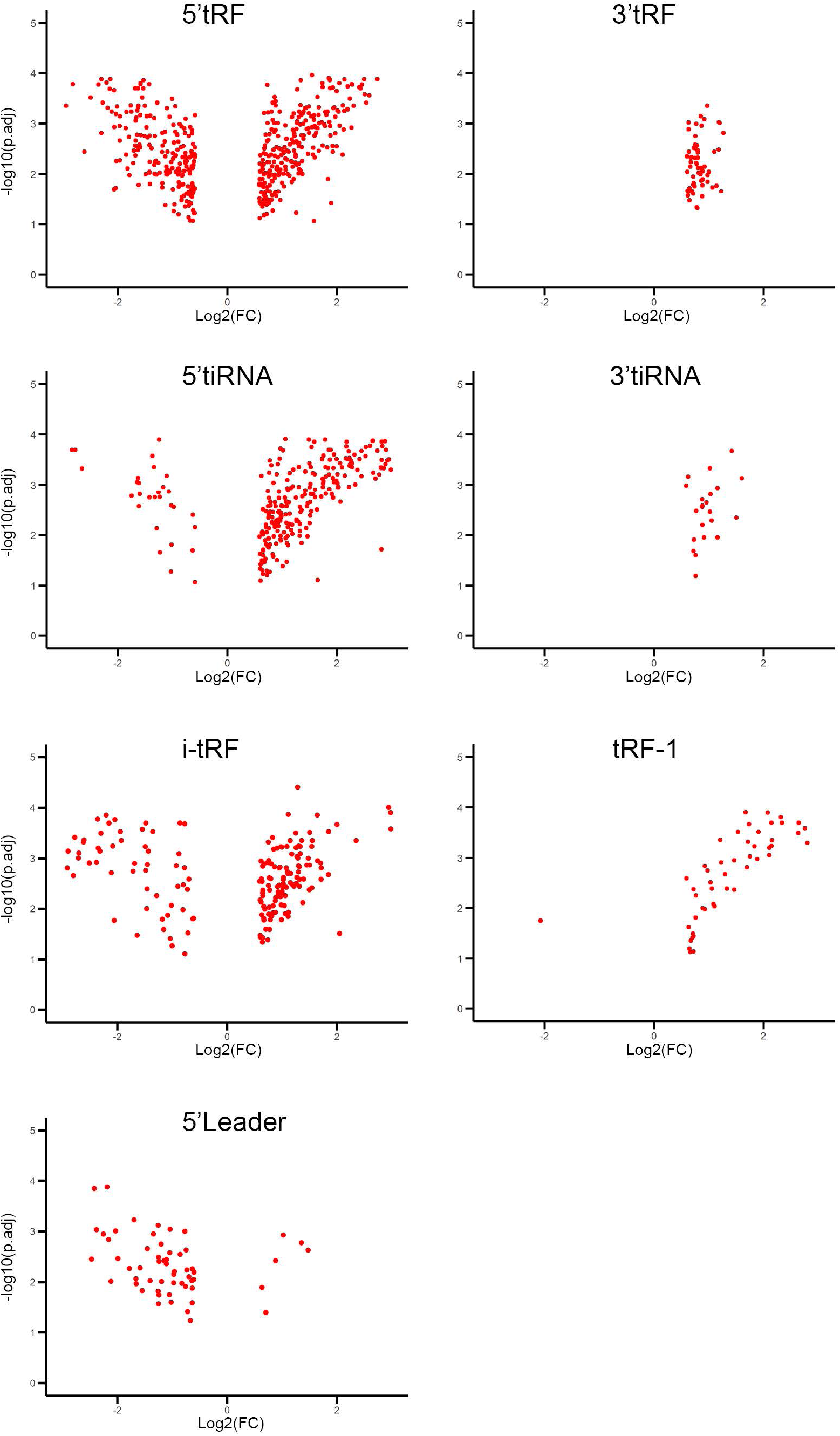
Volcano plots of different classes of tsRNAs that are significantly changed during serum stimulation.

**Figure S3.**
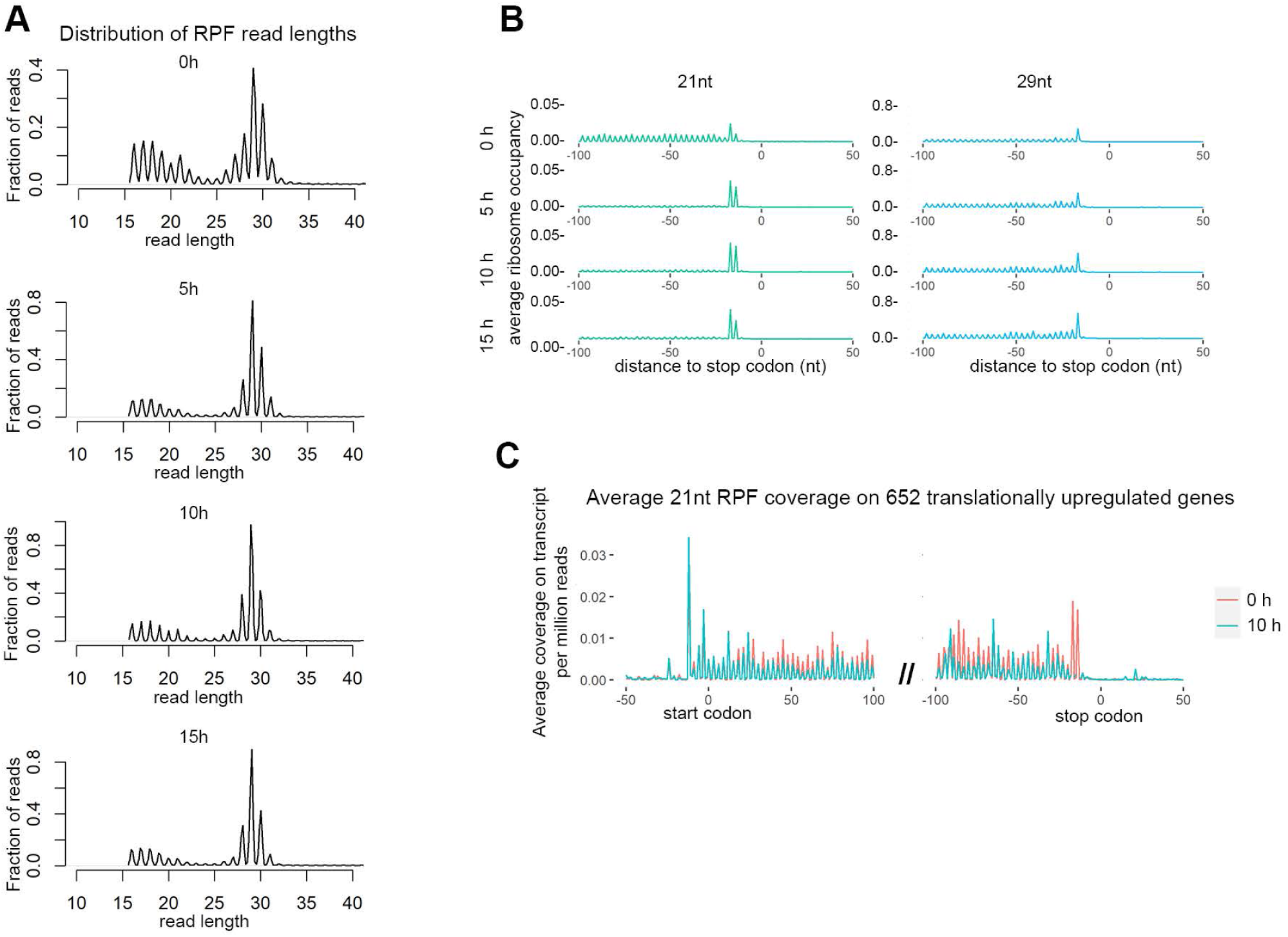
Properties of 21nt and 29nt ribosome protected fragments. (A) Read length distributions of ribosome protected fragments (RPF) from serum starved cells (0 hr) and serum stimulated cells (5 hr, 10 hr and 15 hr). (B) Average ribosome occupancies aligned at stop codons using 21nt RPFs (left) and 29nt RPFs (right). (C) Ratio of 21/29nt RPFs utilized at each codon for 0h, 5h 10h and 15h (aligned by P-site). (D-E) Average ribosome pausing index at P-site (D) and E-site (E) during serum starvation at each codon. (F) Average ribosome occupancies of 21 nt RPFs for the 652 translationally upregulated genes at 0 h and 10 hr.

**Figure S4.**
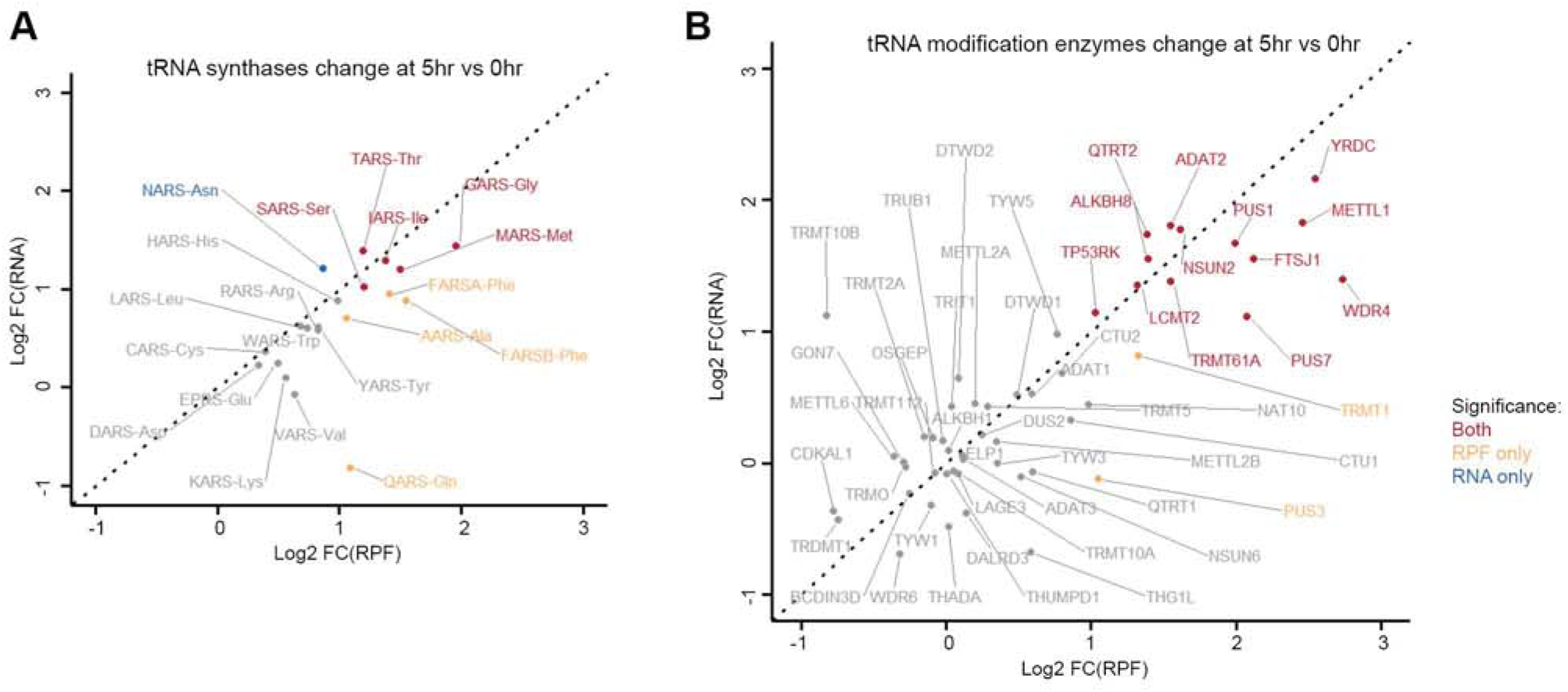
Many aminoacyl-tRNA synthetases and tRNA modifiers are upregulated by serum. (A-B) Log2 fold changes of RNA vs RPF abundance between 5hr and 0hr for aminoacyl-tRNA synthetase genes (A) and tRNA modifier genes (B).

**Figure S5.**
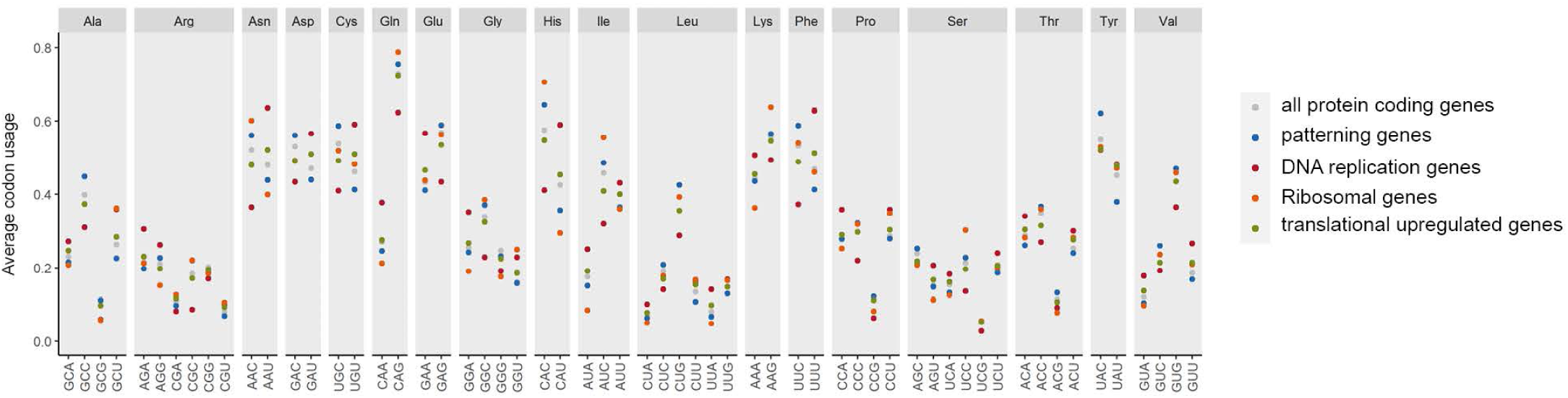
Average codon usage by protein coding genes, patterning genes, and DNA replication genes from the siRNA screen, ribosomal genes, and the 652 translationally upregulated genes.

**Figure S6.**
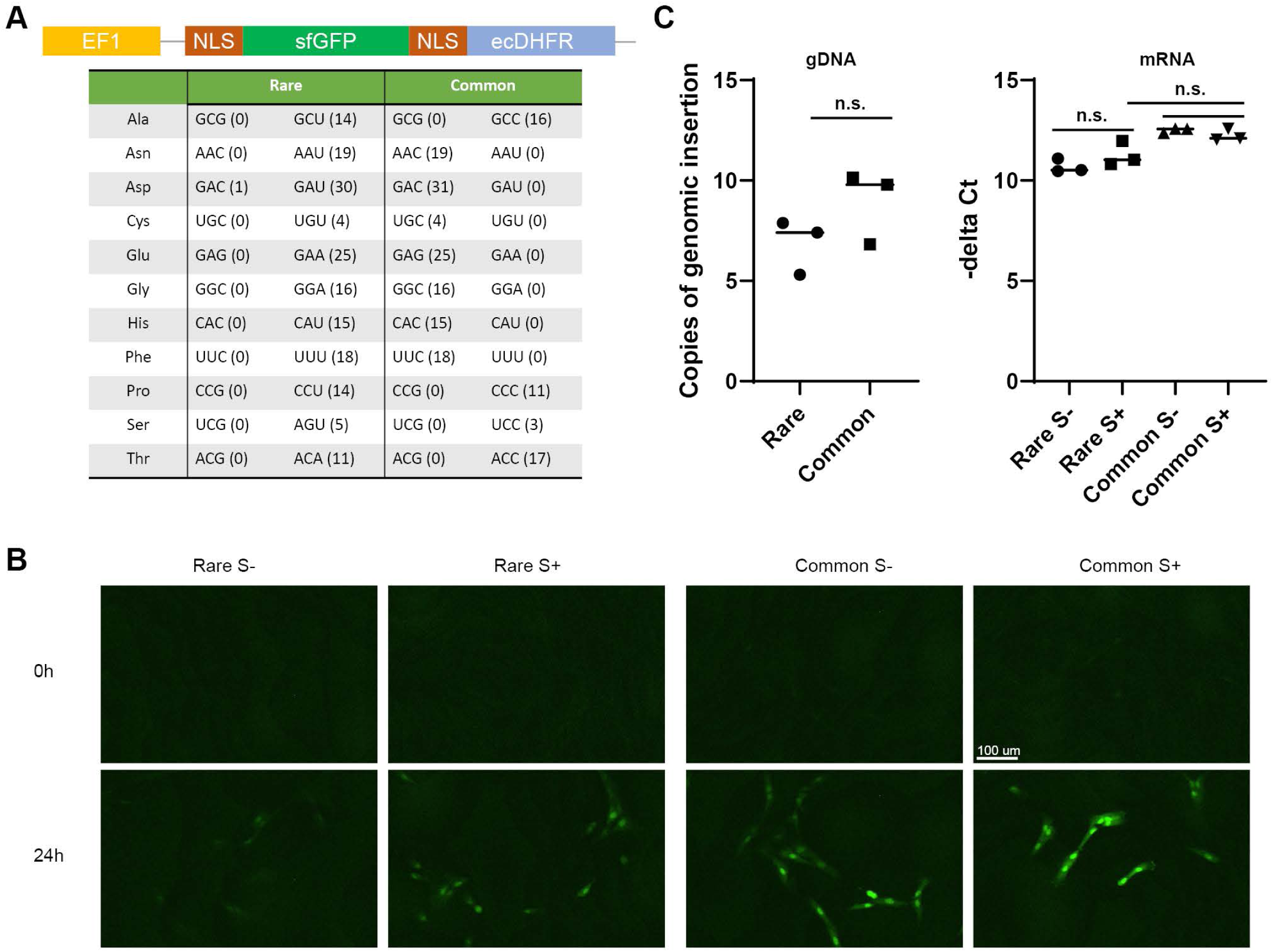
Fluorescent reporters detect the impact of codon usage during the serum response. (A) Codon changes of the NLS-sfGFP-NLS-ecDHFR coding region for “Rare” codon-containing and “Common” codon-containing reporters. Codons substituted into the “Rare” reporter are those preferentially present in our set of translationally regulated DNA replication genes, which are relatively rare in other gene sets including all genes. Codons substituted into the “Common” reporter are those most highly represented in the human genome overall. (B) Fluorescent images of Rare and Common reporters under serum starvation or stimulation for 0h and 24h. (C) qPCR measurements of codon reporter gene copies inserted into genomic DNA (left) and mRNA expression levels during serum starvation and stimulation (right). p-values were determined by unpaired two-tailed t-test; ****< 0.0001, **< 0.01, *< 0.05.

